# The role of superstition of cognitive control during neurofeedback training

**DOI:** 10.1101/2021.09.14.460252

**Authors:** Doris Grössinger, Florian Ph.S Fischmeister, Matthias Witte, Karl Koschutnig, Manuel Ninaus, Christa Neuper, Silvia Erika Kober, Guilherme Wood

## Abstract

**Background:** Real-time fMRI neurofeedback is growing in reputation as a means to alter brain activity patterns and alleviate psychiatric symptoms. Activity in ventral striatum structures is considered an index of training efficacy. fMRI response in these brain regions indicates neurofeedback-driven associative learning. Here we investigated the impact of mere superstition of control as observed during neurofeedback training on patterns of fMRI activation.

**Methods:** We examined the brain activations of a large sample of young participants (n = 97, 50 female, age range 18-54yrs) in a simple fMRI task. Participants saw a display similar to that typically used for real-time fMRI. They were instructed to watch the bars’ movements or to control them with their own brain activity. Bar movements were not connected with brain activity of participants in any way and perceptions of control were superstitious. After the pretended control condition, they rated how well they were able to control the bars’ movements.

**Results:** Strong activation in the basal ganglia and ventral striatum as well as in large portions of the anterior insula, supplementary motor area, and the middle frontal gyrus due to the superstition of brain control.

**Conclusions:** The superstition of control over one’s own brain activity in a pretended neurofeedback training session activates the same neural networks as neurofeedback-driven learning. Therefore, activity in the basal ganglia and ventral striatum cannot be taken as evidence for neurofeedback-driven associative learning unless its effects are proven to supersede those elicited by appropriate sham conditions.

## Introduction

Throughout recent years real-time fMRI neurofeedback has grown from a purely scientific application enabling healthy controls to gain control over well-localized brain areas (1) into a viable clinical tool to treat various neurological and psychiatric disorders. Today RT-fMRI neurofeedback finds its application in treating a wide variety of clinical populations see, e.g., Tursic et al. (2) or Dudek et al. (3) for general reviews. For more specific applications, compare Patel et al. (4) for a recent meta-analysis on treating chronic pain, Trambaiolli et al. (5) for a systematic review on neurofeedback treatment in major depression, Pimenta et al. (6) for the treatment of ADHS, or Wang et al. (7) for the potential of RT-fMRI neurofeedback in stroke rehabilitation.

In all these applications, the BOLD-response in specific brain regions or in more complex networks is extracted in real-time and fed back to the participant. This way, patients can use the feedback information to learn how to regulate the activation levels in specific areas or networks and thereby improve symptoms (8).

Following initial controversies, the general framework for RT-fMRI neurofeedback studies is well defined, c.f Heunis et al. (9), Sorger et al. (10), or Fede et al. (11) for reviews and recommendations on design and methods. Contrary to this, the mechanisms by which neurofeedback works are still controversial. While there is a general agreement on a multiple component model driving the effects of neurofeedback (12), the contribution of individual components is not reported consistently, and still requires specific evaluation. This includes among others the influence of pretraining brain activity (13), target-independent neuronal mechanism (14), psychological factors like attention, motivation, and personality factors (15), or even the role and training of executive functioning (16, 17). Among all these influencing factors, cognitive control (18) and the influence of superstition of control on one’s own brain activation has shown to be an essential factor during neurofeedback (19, 20). Particular protocols need to be employed to investigate and to control for the influence of cognitive control. However, such protocols are rarely employed since they involve an extra sham condition which induces the believe of receiving real feedback about their own brain activity and well-defined brain regions where possible effects of cognitive control are expected (21, 22).

A meta-analysis of fMRI neurofeedback studies in healthy participants (14) revealed a network of brain regions engaged in mediating neuronal self-regulation during neurofeedback. The authors found systematic activation in the ventrolateral and dorsolateral prefrontal cortex, the anterior cingulate, the temporoparietal area, visual cortex, including the temporo-occipital junction and the anterior insula, bilaterally. Moreover, analyses revealed activation in the basal ganglia and the thalamus. Interestingly, these brain regions were found to be activated independently of the targeted region-of-interest, i.e., irrespective of the nature of the feedback task.

The interpretation of the activation in the basal ganglia and the ventral striatum is generally considered straightforward in fMRI neurofeedback studies. Activation in the ventral striatum reflects foremost reward representation (23). These regions are typically activated during associative learning (24), such as operant conditioning or reinforcement learning. Successful self-regulation of brain signals thus occurs when desired brain responses are reinforced by contingent feedback and/or reward (8). This view is supported by animal studies where neurofeedback learning was disturbed by blocking NMDA receptors in the basal ganglia (8, 25, 26).

Furthermore, studies in rodents showed that neurofeedback learning activates cortical–basal ganglia loops responsible for procedural learning (26). Associated activation in the ventral striatum is here seen as an unconscious reward process underlying the procedural and implicit nature of neurofeedback learning (8, 26). Accordingly, the ventral striatum is activated during fMRI neurofeedback even without awareness (27). Paret et al. (28), targeting the insula, observed correlations between the levels of activation in the ventral striatum and the anterior insula. This activation in the striatum was interpreted as a function of how well participants were able to modulate the activation level in the anterior insula in the desired direction (up or down in two different groups). Based on these prior findings, activation in the basal ganglia and ventral striatum structures are considered as indicators of neurofeedback-driven associative learning.

However, this evidence only shows that effective neurofeedback activates the basal ganglia and the ventral striatum, but this activation cannot be distinguished from the superstition of control (19, 21, 22) over one’s brain activity. To achieve such a distinction a placebo condition must be included in the study design (29, 30).

The aim of the present study was to investigate the influence of varying levels of superstition of control on the activation observed in the basal ganglia and ventral striatum during fMRI neurofeedback. To this end, we used a sham neurofeedback task, which induced participants to believe in receiving real feedback about their brain activity. This task consisted of three conditions, or levels of control: “*monitor static bars*” for watching static bars, “*monitor moving bars*” for watching moving bars pretended to be participants’ brain activity and “*control moving bar*” for trying to control the bars and keep the bar high up as long as they could. Using specific and distinct comparisons these three conditions will allow disentangling reward from superstitious control over reward and will help to characterize the role of the latter in neurofeedback studies.

## Methods and Materials

### Participants

Ninety-seven healthy adults (50 female; mean age = 27.6, SD = 7 years, range 19 to 54 years) participated in the current study following written informed consent. Three participants were excluded from further analyses because of bad fMRI data quality. All participants were naïve to the purpose and aims of the study and had never taken part in neurofeedback training before. Inclusion criteria were normal or corrected to normal vision and absence of any major psychiatric or neurological disorder. Participants were recruited via email and word of mouth; selection criteria were verified on-site using demographic questionnaires. All procedures were approved by the local ethics committee (University of Graz) in accordance with the Declaration of Helsinki. Participants were informed that they could cease participation at any point in time without further consequences or do not have to provide any justification.

### Experimental design

The task employed in the present study was comparable to procedures adopted during real-time fMRI neurofeedback training, except that the feedback presented to participants was sham (19). Participants were told that their brain activity was directly coupled to the bar movements and that in some conditions, they should control position of the bar while not moving their body or eyes. In this active experimental condition, “*control moving bars*”, participants were asked to increase the middle bar presented on the visual display while keeping the two outer bars low. In the two other conditions, participants were asked to monitor the bars only. In the high-level control condition “*monitor moving bars*”, participants were asked not to interfere with the middle bar movements but to monitor them passively. In the low-level control condition “*monitor static bars*”, participants were asked again just to observe the bars, which did not move (Figure 1).

**Figure 1.**
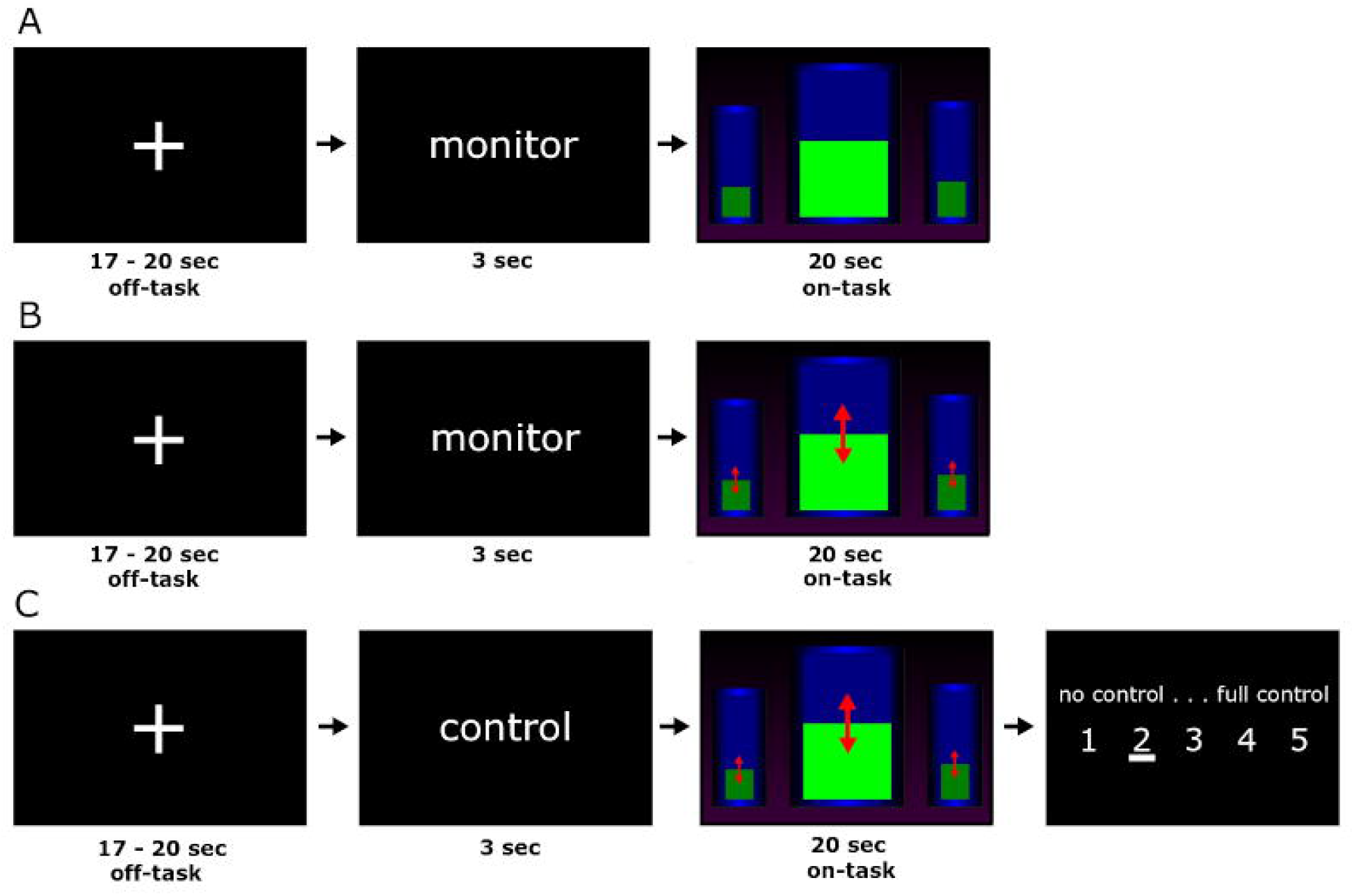
(A) Example of a complete trial of the “monitor static bar” condition; (B) Example of a complete trial of the “monitor moving bar” condition; (C) Example of a complete trial of the “control moving bar” condition (the red arrows represent the possible directions of bar movement). The on-task and off-task conditions are labeled explicitly.

Each trial started with a cross-hair presented for about 18.5s (random jitter from 17 to 20s). Participants then received the instructional cue words “control” or “monitor” for another 3s before the bar display was visible. The feedback-bar was presented for 20s. Immediately following the active control task, participants were asked to rate their level of control on a 5-point rating scale with the anchor words “no control” and “full control”. Each condition was repeated five times, the order of presentation of the three experimental conditions was randomized per subject.

Participants were instructed that there was no explicit strategy on how to control the bars, except for the broad instruction to relax the body, not to move, and to keep focused on the bar. These instructions were identical to regular neurofeedback training of the sensorimotor rhythm using EEG (31-33). The bars’ movements representing neurofeedback were obtained from an actual but unrelated EEG neurofeedback training session. This session was first artifact corrected, cut into 20-second intervals, inspected for signal jumps (affected intervals were excluded), and then arranged in four different ways (forward, backward, forward split into four sub-blocks and re-ordered, backward split into four sub-blocks and re-ordered) yielding four different versions of the task. These task versions were smoothed with a 1s moving average to prevent jumps, particularly in the split versions. Each participant was presented with one of these four generated versions at random. Only after the completion of data collection of all participants, the sham design was disclosed to participants.

### MRI image recording

Images were acquired with a 3.0 Tesla Skyra MRI scanner (Siemens, Erlangen Germany) at the MRI-Lab Graz (Austria) using a 32-channel head coil. Anatomical images were collected using a T1-weighted MPRAGE sequence (TR = 1650ms; TE = 1.82ms; flip-angle = 8°; slice thickness = 1mm; 256 × 256 × 192 acquisition matrix; voxel dimensions = 1 × 1 × 1mm; TI = 1000ms). Functional images were acquired using a T2* weighted multiband gradient-echo pulse imaging sequence (TR = 1250ms; TE = 40ms, flip angle = 60°; slice thickness = 2.5mm; 96 × 96 matrix; voxel dimensions = 2.5 × 2.5 × 2.5mm), providing whole-brain coverage in 52 slices. Visual stimuli were presented using an MR-compatible monitor visible via a mirror attached to the head coil. Ratings of the perceived level of control were recorded via an MR-compatible response box. Buttons under participants’ index and ring fingers were used to navigate through the rating scores mentioned above; the button under their middle finger served to confirm their choice. Because participants were required to press the buttons well after the 20s task block and without time constraints, leakage of preparatory movement-related activations into the cognitive demanding self-regulation period was precluded.

### MRI image preprocessing

Neuroimaging data were processed in Matlab 2016a (The MathWorks, Natick, MA) and SPM12 (http://www.fil.ion.ucl.ac.uk/spm/). We used the DPARSFA toolbox (http://www.restfmri.net) for image preprocessing. First, fMRI data were slice time corrected using the middle slice as reference and then realigned to the first volume. Next, images were coregistered to the T1 image and spatially normalized to the Montreal Neurological Institute (MNI) as implemented in SPM12. After spatial transformation of the realigned T2* volumes, images were smoothed with an 8-mm FWHM Gaussian kernel to improve group-level statistics.

### Statistical analysis

Behavioral data were obtained following each block of the “*control moving bars*” condition, indicating the subjective estimation of neurofeedback control achieved during each run. Both, the average rating as well as its variability were considered as good indicators of the overall degree of conviction to control the bar movements with one’s own brain activity. As described above, these ratings ranged from 1 to 5, indicating no control at all (rating = 1), moderate levels of control (rating = 3), or a high subjective level of control (rating = 5). These ratings obtained during the “*control moving bars*” condition were analyzed using an ANOVA model with a between-subjects factor representing task version (4 levels) and a within-subjects factor representing time (5 levels, since this control condition was presented five times per participant). This ANOVA was conducted to investigate possible differences induced by the four versions of the presented feedback on the participant’s belief of control.

Functional imaging data were analyzed using SPM12. Signals were modeled at the first level using a block design. For this individual analyses two separate models were generated, each with all events defined explicitly (i.e. fixation cross, instructional cue, “feedback”, and rating) and convolved with the HRF. For the first model, serving as replication to Ninaus et al. (19), the “feedback” event was subdivided into the three conditions, i.e., the different general levels of control employed by the participants (“*monitor static bars”* for watching static bars, “*monitor moving bars*” for watching moving bars and “*control moving bars*” for trying to control the bars). For the second model, we were interested in neuronal activity accounting for the individual experience of control. To this aim, the “feedback” event was modeled using parametric modulators, one defining the presented level of control over the stimulus (the condition) and the other representing the subjective ratings given by the participant for every single event. The first modulator, termed “*levels of control*”, aims at catching all the information associated with different presentations of the feedback and the participants’ general belief to control the presented bars actively. Here, the “*watch static bars*” condition was included in this modulator to account for static visual representations of no interest. The second modulator, termed “*rating*” aims at representing the individual belief of control over the moving bars ranging from 1 (no control) to 5 (full control); for the two monitoring conditions, “static bar” and “moving bar”, this modulator was zeroed for consistency. The six realignment parameters of the motion correction procedure (translation and rotation) were entered as covariates of no interest into the first-level analyses to remove residual artifacts of head movements.

Single-subject contrasts obtained from first-level modeling were then brought to the group level. To replicate previous research, we calculated a within-subjects t-test comparing “*control moving bars*” > “*watch moving bars*”. both obtained from the classical first-level model. For the following analyses, the second parametric modulator model was used as input. Activation corresponding to participants’ general belief to actively control the presented bars was captured by calculating a one-sample t-test for the parametric modulator representing “*levels of control”*. Activations related to the subjective belief of control over the presented bar were analyzed by calculating a one-sample t-test for the parametric modulator representing individual ratings. Since both modulators are orthogonal to each other, observed activation can be considered non-overlapping.

Whole-brain analysis results were family-wise error (FWE) corrected for multiple comparisons on the cluster-level employing a threshold of p < 0.05, with a minimum cluster size of 20 voxels. All results are masked for gray matter of the brain and the cerebellum.

## Results

### Behavioral ratings

During the debriefing, all participants rated perceived success in controlling the bars when instructed to do so. The mean ratings obtained following the “*control moving bars*” condition across all participants was 3 (M = 3.15, SD = 0.89; Range: 1-5). Three out of all 97 participants, including those subjects excluded due to low fMRI data quality, showed no variance in their ratings (two participants gave a continuous rating of 1 and one a continuous rating of 2). Ratings did not differ significantly across task versions (F(3, 90) = 0.57, p > 0.05).

### fMRI Activation

For the whole-brain analysis we evaluated the three contrasts described before: the parametric modulators representing “*levels of control*” and “*rating*”, and the contrast “*control moving bars*” versus “*watch moving bars*”. The activation maps for these contrasts are shown in Figures 2-4; corresponding t-values, MNI space coordinates, and number of significant voxels can be found in Table 1-3.

**Table 1.**
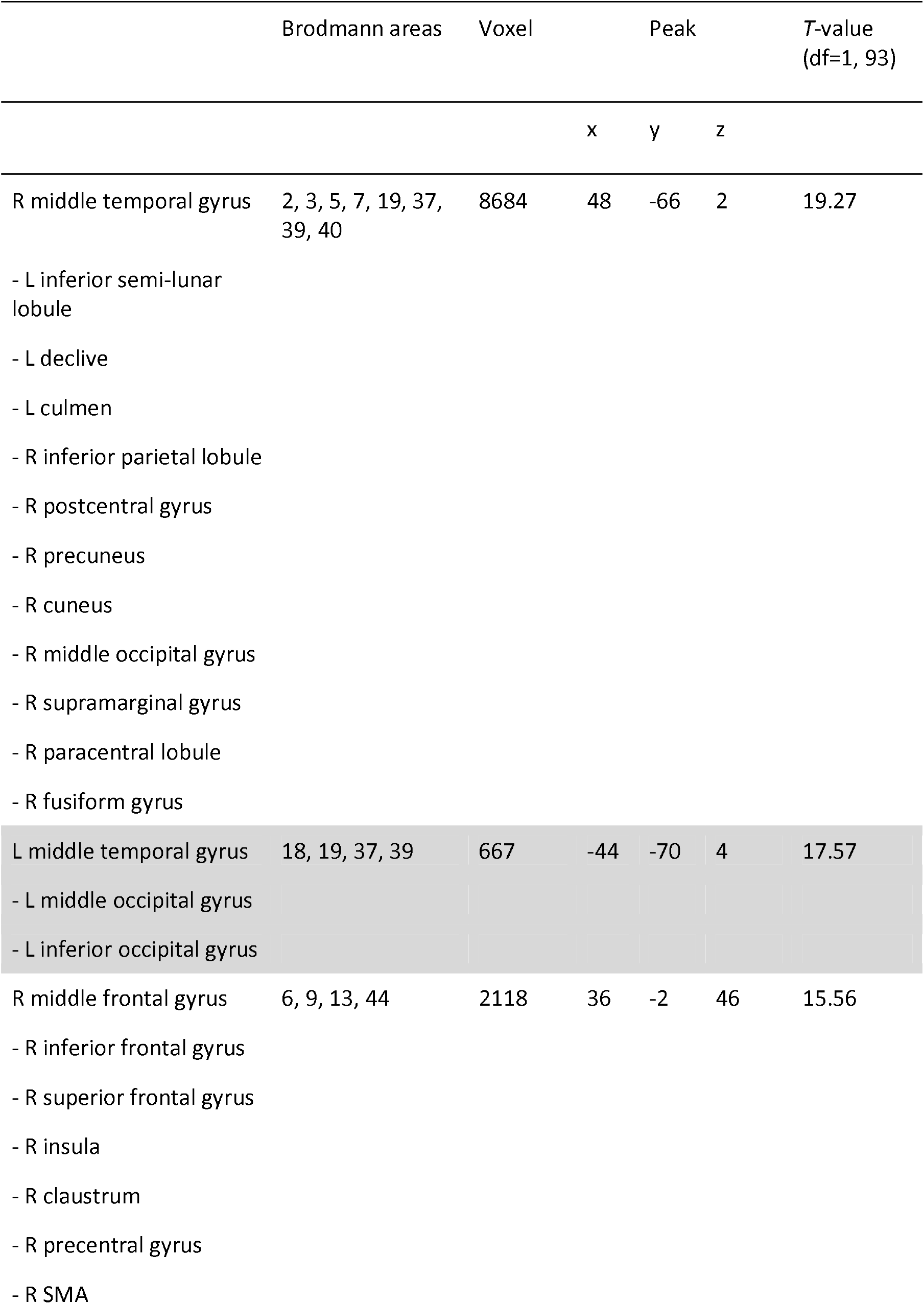
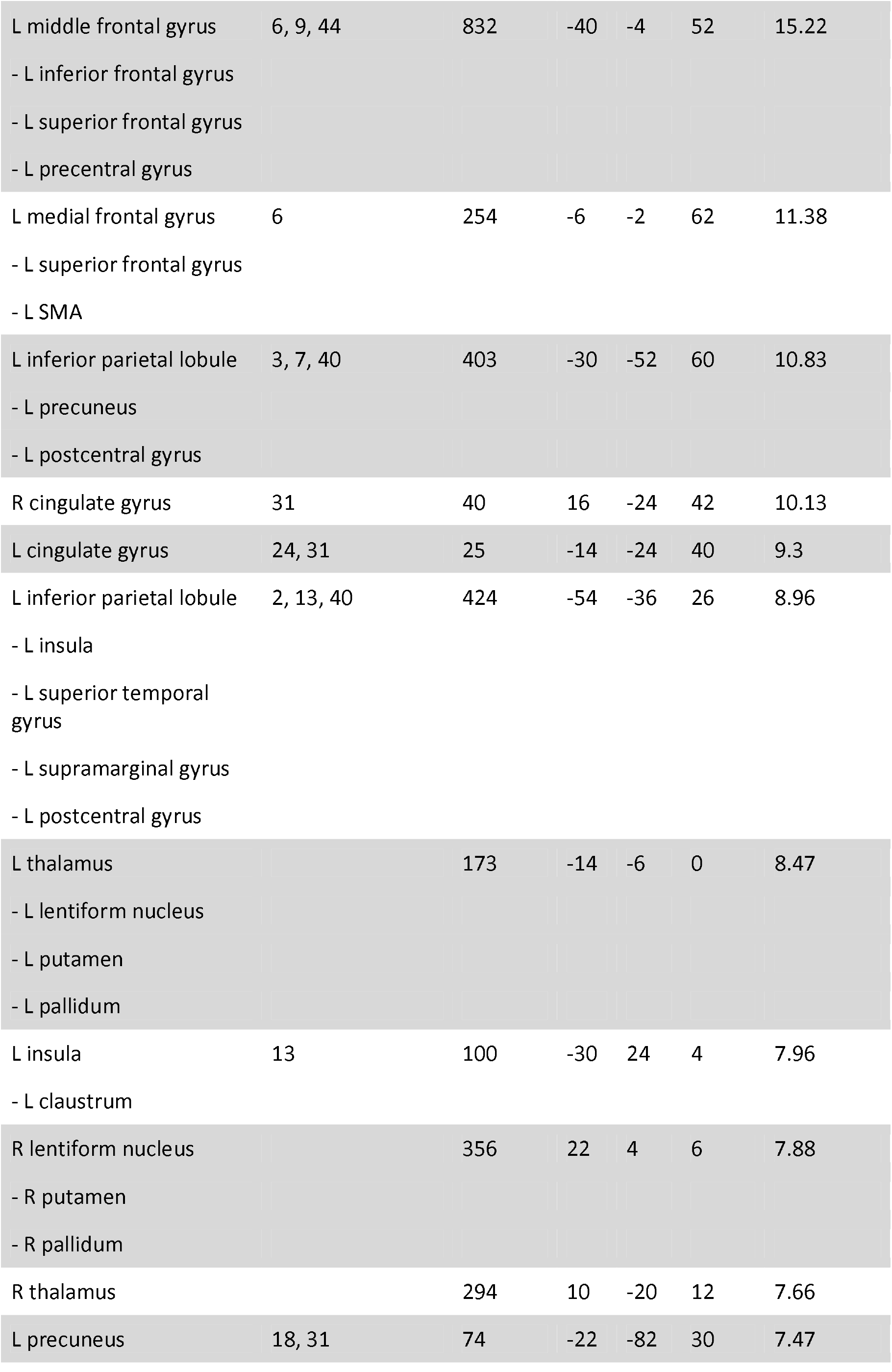
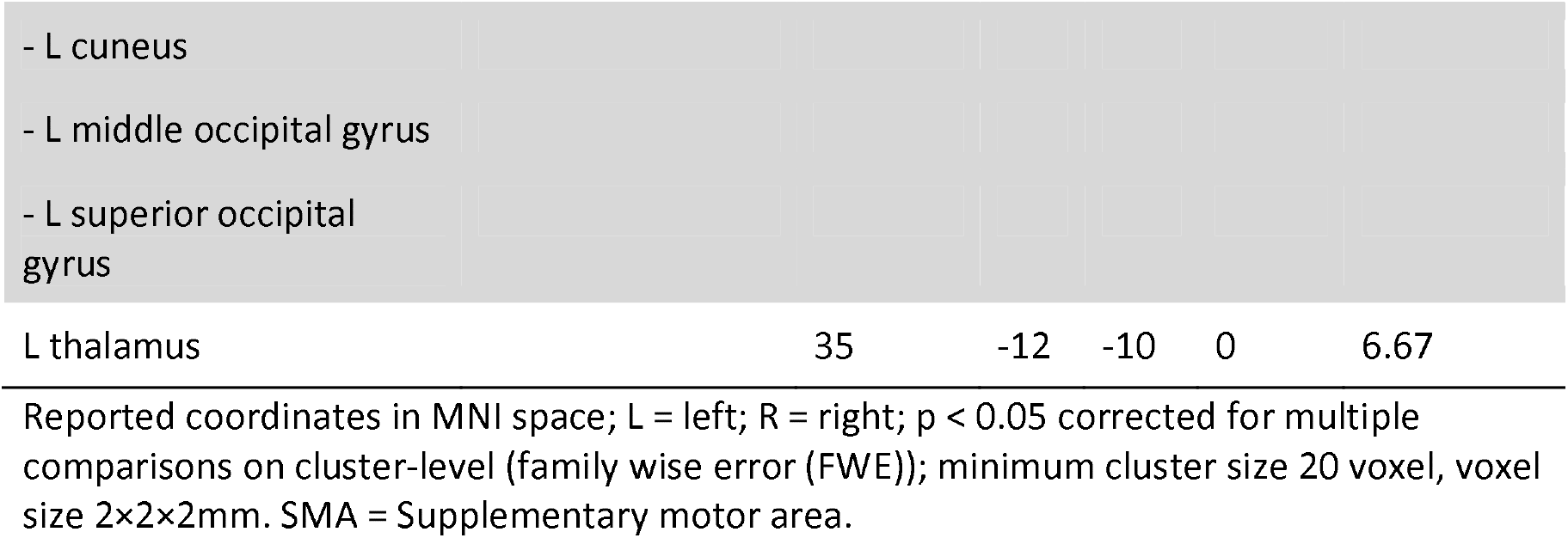
Brain regions activated for “levels of control”

**Figure 2.**
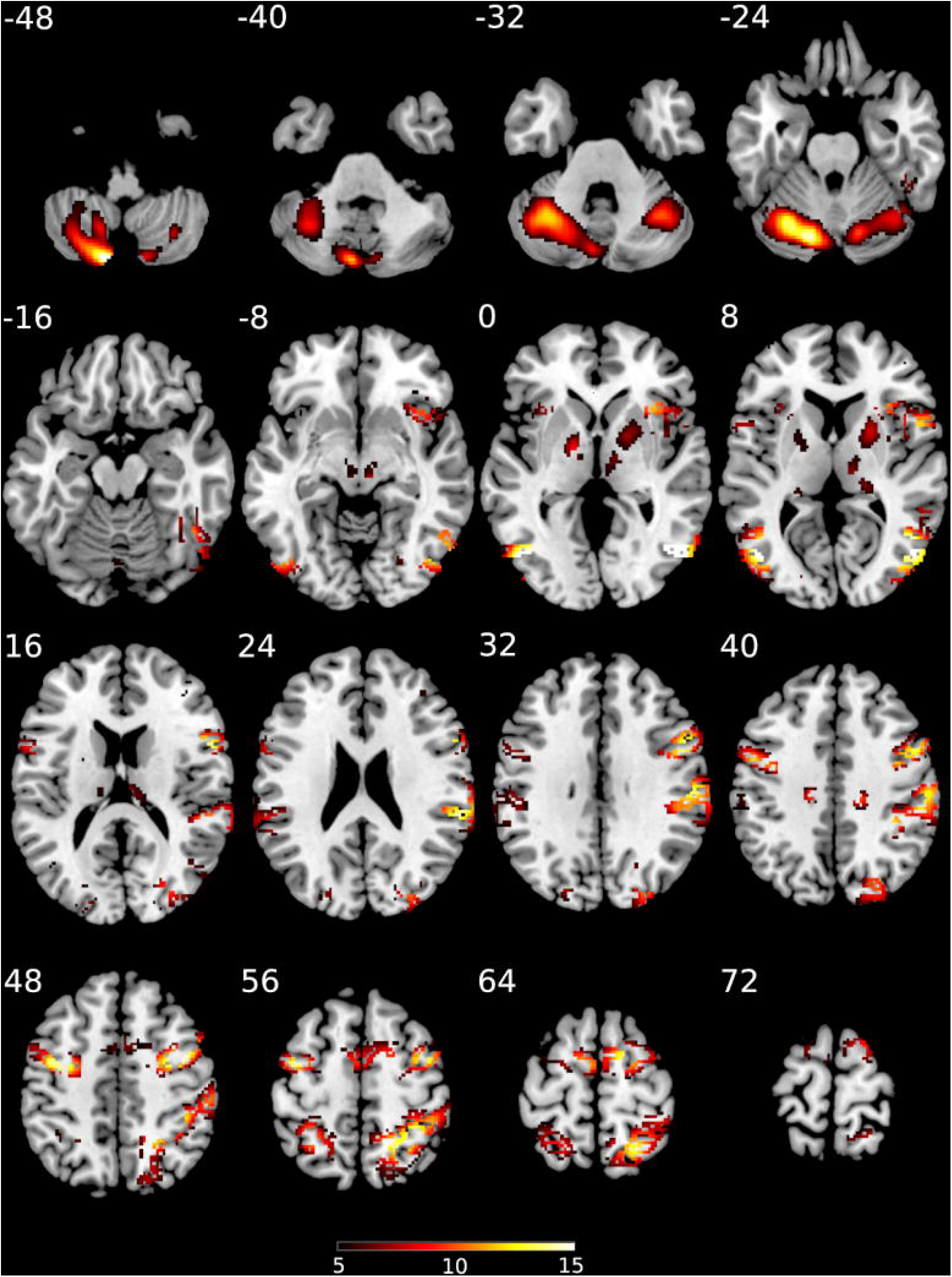
t-score map for activation “levels of control”; p < 0.05 corrected for multiple comparisons on cluster-level (family wise error (FWE)); minimum cluster size = 20 voxel; voxel size = 2×2×2mm; the colors represent the respective T-statistics; MNI coordinates; transversal slices; numbers in the top left corners are the respective slice coordinates.

The aim of the modulator “*levels of control*” was to show activation related to participants’ general belief to actively control the presented bars. Significant activation related to this general belief was found within the thalamus, the basal ganglia, the ventral striatum, cerebellum, insula, cingulate gyrus, inferior parietal lobule, as well as within the pre- and postcentral gyrus, and the supplementary motor area (SMA). Further significant clusters were observed within parts of the frontal, temporal, and occipital cortex (c.f. Table 1 and Figure 2). Additionally, we calculated the contrast “*control moving bars*” > “*watch moving bars*” for a more specific analysis on regions activated when participants are supposed to control the moving bars versus just watching them. Results for this comparison were similar to our previous contrast except for an markedly increased activation in the striatum and the basal ganglia. This distinction was complimented by different activation patterns in the visual cortex (c.f. Table 2 and Figure 3).

**Table 2.** Brain regions activated for the contrast “control moving bars” > “monitor moving bars”.

|  | Brodmann areas | Voxel | Peak |  |  | T-value<br>(df=1, 186) |
| --- | --- | --- | --- | --- | --- | --- |
|  |  |  | x | y | z |  |
| L + R declive |  | 8866 | 30 | -60 | -28 | 17.12 |
| - R cerebellar tonsil |  |  |  |  |  |  |
| - L + R culmen |  |  |  |  |  |  |
| - L cerebellar tonsil |  |  |  |  |  |  |
| - L + R inferior semi-lunar lobule |  |  |  |  |  |  |
| - L dentate |  |  |  |  |  |  |
| - R nodule |  |  |  |  |  |  |
| - R uvula of vermis |  |  |  |  |  |  |
| L medial frontal gyrus | 6, 13, 44 | 3281 | -2 | -2 | 56 | 15.27 |
| - L SMA |  |  |  |  |  |  |
| - L insula |  |  |  |  |  |  |
| - L inferior frontal gyrus |  |  |  |  |  |  |
| - L precentral gyrus |  |  |  |  |  |  |
| - L lentiform nucleus |  |  |  |  |  |  |
| - L medial globus pallidus |  |  |  |  |  |  |
| - L claustrum |  |  |  |  |  |  |
| - L red nucleus |  |  |  |  |  |  |
| R medial frontal gyrus | 6, 24, 44 | 1409 | 2 | -4 | 56 | 13.65 |
| - R SMA |  |  |  |  |  |  |
| - R cingulate gyrus |  |  |  |  |  |  |
| - R inferior frontal gyrus |  |  |  |  |  |  |
| - R middle frontal gyrus |  |  |  |  |  |  |
| - R precentral gyrus |  |  |  |  |  |  |
| - R superior frontal gyrus |  |  |  |  |  |  |
| R insula | 13, 44 | 1267 | 44 | 4 | 8 | 12.51 |
| - R lentiform nucleus |  |  |  |  |  |  |
| - R medial globus pallidus |  |  |  |  |  |  |
| - R putamen |  |  |  |  |  |  |
| - R claustrum |  |  |  |  |  |  |
| - R precentral gyrus |  |  |  |  |  |  |
| - R thalamus |  |  |  |  |  |  |
| - R superior temporal gyrus |  |  |  |  |  |  |
| - R caudate |  |  |  |  |  |  |
| R red nucleus |  | 622 | 8 | -22 | -8 | 9.35 |
| - R thalamus |  |  |  |  |  |  |
| - R ventral lateral nucleus |  |  |  |  |  |  |
| - R substantia nigra |  |  |  |  |  |  |
| - R caudate |  |  |  |  |  |  |
| L cuneus | 18, 19 | 256 | -8 | -96 | 22 | 7.78 |
| - L middle occipital gyrus |  |  |  |  |  |  |
| R middle frontal gyrus | 9, 10 | 24 | 32 | 42 | 26 | 6.84 |
| - R superior frontal gyrus |  |  |  |  |  |  |
| L postcentral gyrus | 40 | 28 | -64 | -22 | 22 | 6.76 |
| - L inferior parietal lobule |  |  |  |  |  |  |
| R cuneus | 18 | 170 | 14 | -94 | 28 | 6.58 |
| L inferior parietal lobule | 40 | 81 | -56 | -32 | 26 | 6.30 |
| L precuneus | 7 | 38 | -18 | -52 | 62 | 5.83 |
| R precuneus | 7 | 45 | 20 | -50 | 62 | 5.68 |

Reported coordinates in MNI space; L = left; R = right; $p < 0.05$ corrected for multiple comparisons on cluster-level (family wise error (FWE)); minimum cluster size 20 voxel, voxel size 2×2×2mm. SMA = Supplementary motor area.

**Figure 3.**
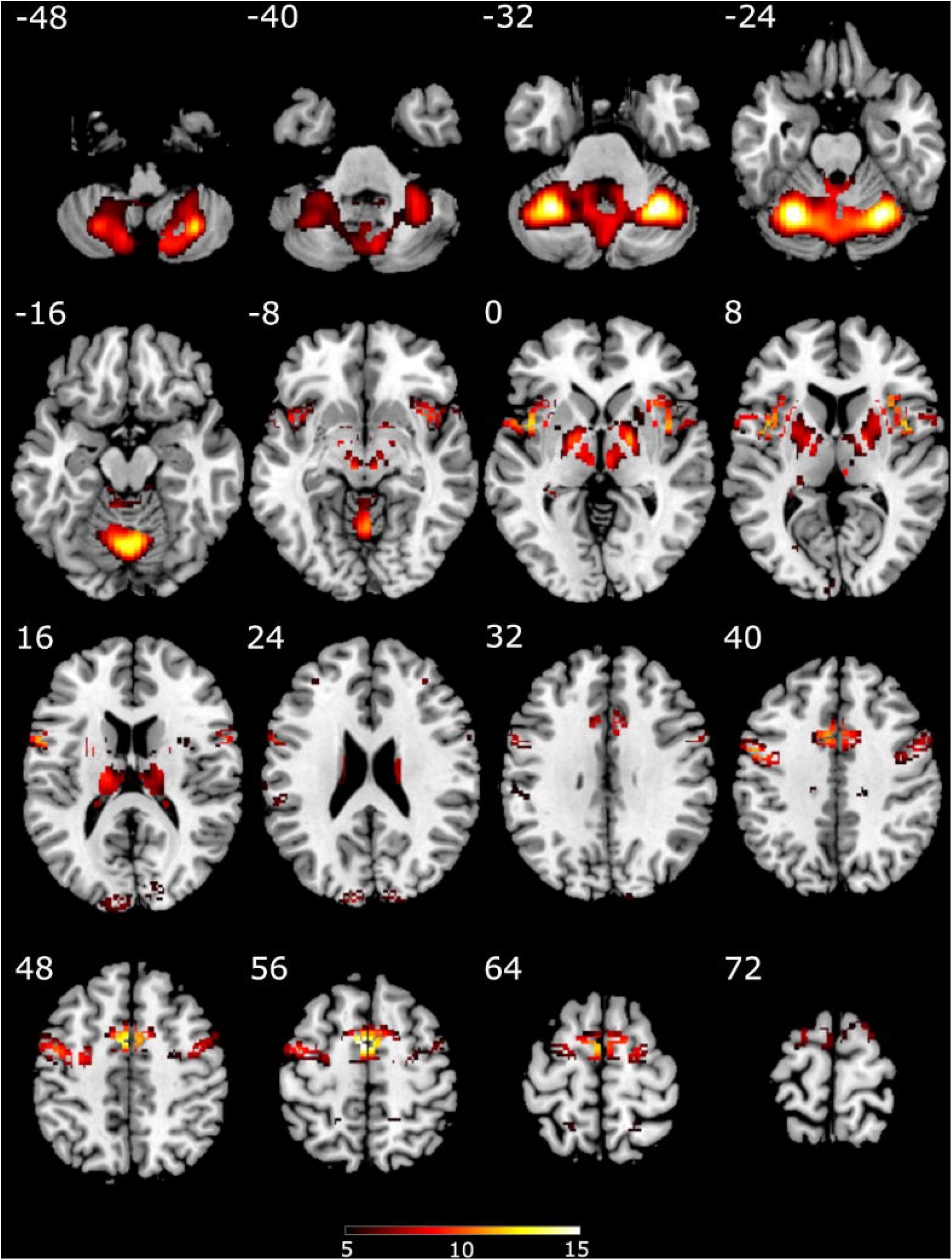
t-score map for activation for the contrast “control moving bars” > “watch moving bars”; p < 0.05 corrected for multiple comparisons on cluster-level (family wise error (FWE)); minimum cluster size = 20 voxel; voxel size = 2×2×2mm; the colors represent the respective T-statistics; MNI coordinates; transversal slices; numbers in the top left corners are the respective slice coordinates.

For the modulator “rating”, we explicitly aimed to differentiate between individual levels of belief of control as indicated by the different ratings participants reported after the five control the moving bar conditions. Activations associated with this modulator were comparable to the previous contrast “*control moving bars*” > “*watch moving bars*”. Again, we obtained widespread activations in the thalamus, the basal ganglia, the ventral striatum, cerebellum, insula, cingulate gyrus, inferior parietal lobule, pre- and postcentral gyrus, supplementary motor area (SMA), parts of the frontal and temporal cortex (c.f. Table 3 and Figure 4).

**Table 3.** Brain regions activated for “rating”

|  | Brodmann areas | Voxel | Peak |  |  | T-value<br>(df=1, 93) |
| --- | --- | --- | --- | --- | --- | --- |
|  |  |  | x | y | z |  |
| L + R declive |  | 8438 | -30 | -60 | -26 | 15.31 |
| - L + R culmen |  |  |  |  |  |  |
| - L + R inferior semi-lunar<br>lobule |  |  |  |  |  |  |
| - L fastigium |  |  |  |  |  |  |
| - R uvula |  |  |  |  |  |  |
| - R cerebellar tonsil |  |  |  |  |  |  |
| L medial frontal gyrus | 6, 9, 13, 22, 44 | 3475 | -2 | -2 | 56 | 13.82 |
| - L inferior frontal gyrus |  |  |  |  |  |  |
| - L middle frontal gyrus |  |  |  |  |  |  |
| - L precentral gyrus |  |  |  |  |  |  |
| - L SMA |  |  |  |  |  |  |
| - L lentiform nucleus |  |  |  |  |  |  |
| - L medial globus pallidus |  |  |  |  |  |  |
| - L putamen |  |  |  |  |  |  |
| - L thalamus |  |  |  |  |  |  |
| - L superior temporal<br>gyrus |  |  |  |  |  |  |
| - L insula |  |  |  |  |  |  |
| - L claustrum |  |  |  |  |  |  |
| R superior frontal gyrus | 6, 9, 24, 44 | 1445 | 2 | 2 | 56 | 12.96 |
| - R medial frontal gyrus |  |  |  |  |  |  |
| - R inferior frontal gyrus |  |  |  |  |  |  |
| - R middle frontal gyrus |  |  |  |  |  |  |
| - R SMA |  |  |  |  |  |  |
| - R cingulate gyrus |  |  |  |  |  |  |
| - R precentral gyrus |  |  |  |  |  |  |
| R insula | 6, 13, 22, 44 | 1160 | 44 | 4 | 10 | 11.87 |
| - R lentiform nucleus |  |  |  |  |  |  |
| - R medial globus pallidus |  |  |  |  |  |  |
| - R lateral globus pallidus |  |  |  |  |  |  |
| - R lentiform nucleus |  |  |  |  |  |  |
| - R putamen |  |  |  |  |  |  |
| - R claustrum |  |  |  |  |  |  |
| - R thalamus |  |  |  |  |  |  |
| - R precentral gyrus |  |  |  |  |  |  |
| - R superior temporal gyrus |  |  |  |  |  |  |
| R thalamus |  | 700 | 16 | -8 | 10 | 8.76 |
| - R caudate |  |  |  |  |  |  |
| L inferior parietal lobule | 13, 40 | 193 | -54 | -34 | 24 | 7.95 |
| - L insula |  |  |  |  |  |  |
| - L supramarginal gyrus |  |  |  |  |  |  |
| R inferior parietal lobule | 2, 3, 40 | 129 | 66 | -30 | 28 | 7.1 |
| - R postcentral gyrus |  |  |  |  |  |  |
| - R supramarginal gyrus |  |  |  |  |  |  |
| L cuneus | 18, 19 | 194 | -8 | -96 | 22 | 7.08 |
| - L superior occipital gyrus |  |  |  |  |  |  |
| R cingulate gyrus | 31 | 29 | 16 | -26 | 40 | 6.83 |
| R superior parietal lobule | 7 | 118 | 20 | -50 | 64 | 6.1 |
| - R postcentral gyrus |  |  |  |  |  |  |
| L precuneus | 7 | 45 | -18 | -52 | 62 | 6.33 |
| - R superior parietal lobule |  |  |  |  |  |  |
| R cuneus | 19 | 63 | 16 | -96 | 28 | 5.86 |
| - R superior occipital<br>gyrus |  |  |  |  |  |  |

Reported coordinates in MNI space; L = left; R = right; $p < 0.05$ corrected for multiple comparisons on cluster-level (family wise error (FWE)); minimum cluster size 20 voxel, voxel size 2×2×2mm. SMA = Supplementary motor area.

**Figure 4.**
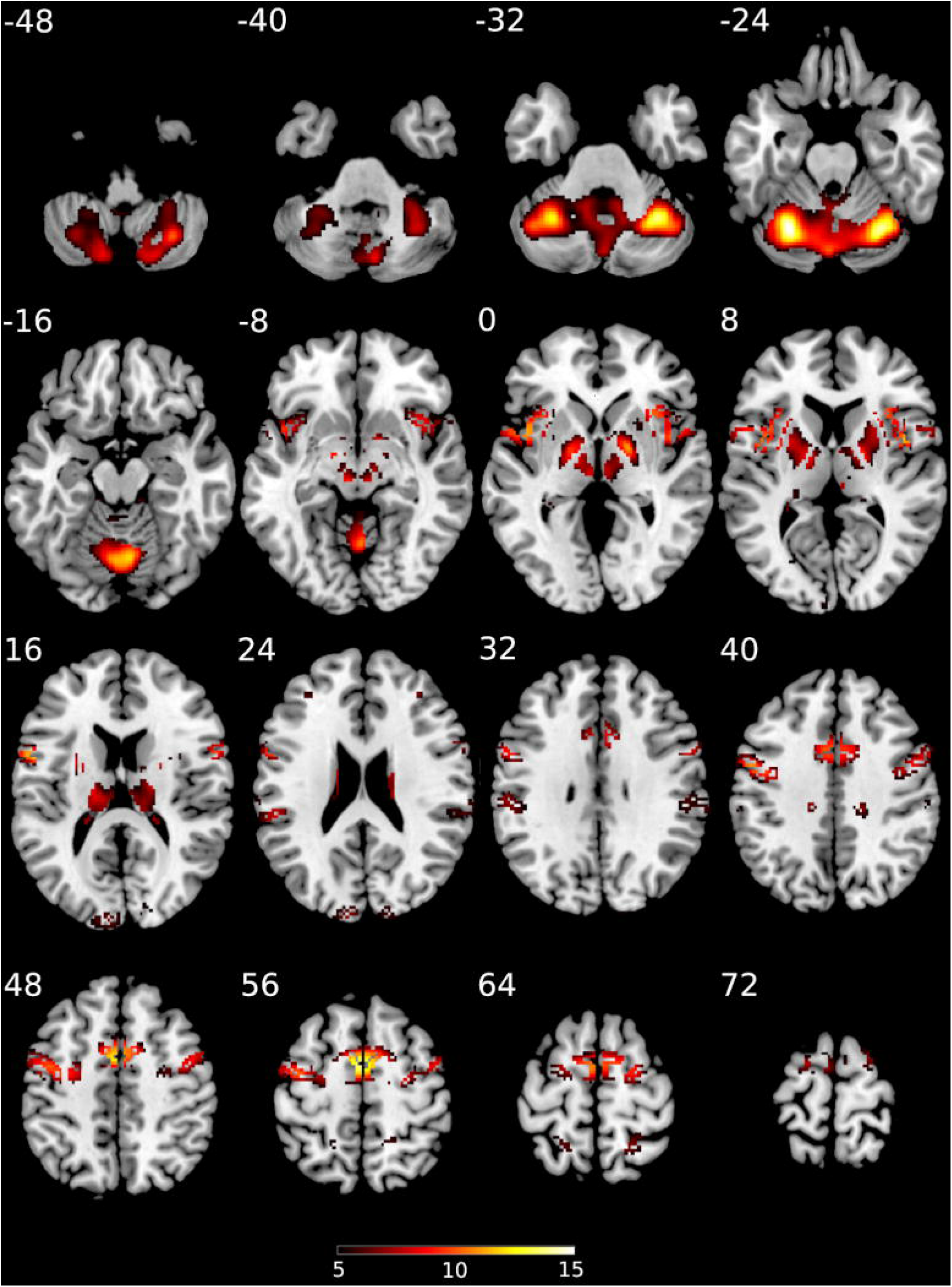
t-score map for activation for participant “rating”; p < 0.05 corrected for multiple comparisons on cluster-level (family wise error (FWE)); minimum cluster size = 20 voxel; voxel size = 2×2×2mm; the colors represent the respective T-statistics; MNI coordinates; transversal slices; numbers in the top left corners are the respective slice coordinates.

## Discussion

Real time fMRI feedback has gained significant popularity not only in the neuroscientific community. One of the most important aspects of effective neurofeedback training is cognitive control. Cognitive control or the absence of control, i.e. the superstition of control,, do underly the procedural and implicit (21, 22) nature of neurofeedback learning, but are hard to distinguish from feedback induced changes in one’s own brain activity (19).

In the present study, we aimed to disentangle the effects of superstition of control from stimulus-induced activation using a sham neurofeedback task. The large majority of participants (n = 94/97, i.e. 97%) showed at least a moderate degree of belief in successful control of the bar using their own brain activation. In line with the original study by Ninaus et al. (19), a network of brain regions including the anterior insula, bilaterally, the middle frontal gyrus, the supplementary motor area, dorsal anterior cingulate, and visual areas of the brain was activated as well as large portions of the basal ganglia, and the ventral striatum. When comparing the activation intensity across the three conditions (“monitor static bars”, “monitor moving bars” and “control moving bars”) using a parametric modulator we found a strong recruitment of the basal ganglia, including the ventral striatum, when participants were expected to control the feedback bar. Additional analysis revealed similar activation when directly comparing the conditions “*control moving bars*” and “*monitor moving bars*”. These results show that the superstition of control over one’s own brain activity activates the basal ganglia and striatal structures in addition to the control network as believed previously (14). Therefore, the interpretation of the ventral striatum activation as a sole index of genuine learning and plasticity induced by neurofeedback seems problematic in fMRI neurofeedback studies not controlled for effects of superstition of control. This will be further elaborated in the following sections.

### Rating control beliefs

The observed high level of control over the sham feedback shows that the level of insight of young and educated participants on the properties of sham fMRI feedback is sufficiently low to be manipulated by simple instructions. This degree of belief in the authenticity of neurofeedback is in line with recent findings on how strong the effects of suggestion in the context of fMRI can be (34). On the one hand, these strong beliefs are indicative that placebo conditions are well accepted by participants during fMRI neurofeedback experiments. On the other hand, this indicates that a strong neural response to placebo conditions should always be expected (30). We also analyzed the brain activations related to the ratings when instructed to control the moving bars. Results show that participants more strongly recruit basal ganglia and the ventral striatum, when the belief of control is higher. This may indicate the reward processing of the arbitrary feedback signal. These findings also reinforce previous findings pointing at the paramount relevance of adding a placebo condition to fMRI neurofeedback designs to be able to claim its efficiency (30). In summary, sham-conditions do not exclude the possibility that a proportion of the observed activation e.g. in the ventral striatum is due to superstition of control and not to feedback related control itself.

### Neurofeedback control

Activation under sham neurofeedback as found in our study strongly resemble areas belonging to the neurofeedback control network as found by Emmert et al. (14). Contrary to Emmert et al. (14) these strong activations within the control network were observed when participants were instructed to actively control a feedback stimulus they never had control of. Importantly the here found brain areas under sham neurofeedback do replicate our previous findings (19) using sham neurofeedback too. Specifically, the contrast “*control moving bars*” > “*watch moving bars*” represents a one on one replication of Ninaus et al. (19). These findings corroborate the view that these networks responsible for different aspects of neurofeedback are recruited even by the illusion of control (19).

For the current study, despite employing sham feedback, we specifically observed activations in regions previously found by Emmert et at. (14) and described in Sitaram et al. (8) as responsible for neurofeedback learning. For these activations it seems that the effort to comply with task instructions and the subjective perception of task control as indicated by the subjective ratings are mobilizing both control and reward networks as genuine fMRI neurofeedback protocols (14).

For the effects of individual control, indicated by the modulator *“levels of control”*, we generally found activations in the same regions. This indicates that the bar movements are rewarding for participants not only by the pure sight of them but rather under the superstition of control over their movement. These results are in line with previous literature pointing out that not only basic learning processes modulate activation in the ventral striatum, but also psychosocial processes such as social learning (21) and expectations generated by sham manipulations (35). Moreover, wrong or superstitious beliefs are apparently able to alter attention and learning as well as the activation of the ventral striatum and basal ganglia (22). More specifically, the “*control moving bar*” condition seems to elicit a stronger form of “neuroenchantment” (34), which increases the rewarding effect of bar movements through the false belief of control.

Interestingly, different properties of neuroanatomical structures of the neurofeedback control network predict the outcomes of neurofeedback training. For instance, Ninaus et al. (20) showed that the volumes in specific brain regions in the frontal cortex and basal ganglia predict the outcome of sensorimotor rhythm EEG neurofeedback training. Moreover, gray matter volumes in the supplementary motor area and left middle frontal gyrus predicted the outcomes of gamma rhythm training (20). Accordingly, Kober et al. (32) reported a positive association between the gray matter volumes in the right insula and inferior frontal gyrus and the outcomes of sensorimotor rhythm neurofeedback training. Finally, Enriquez-Geppert et al. (16) reported an association between the morphology of the midcingulate cortex and the outcomes of frontal-midline theta neurofeedback. Successful EEG neurofeedback training also led to increases in the gray matter volume in brain structures related to neurofeedback control. For instance, EEG neurofeedback of the low beta rhythm (15-18 Hz) seems to increase gray matter volumes in the right middle frontal gyrus, left superior and inferior frontal gyrus, and right thalamus among other structures (36).

Taken together, these studies and the here presented results suggest that the neurofeedback control network influences the outcomes but is also influenced by the neurofeedback training itself. Therefore, it is not correct to consider that their levels of activation have no direct consequence for neurofeedback learning, although they are also under the influence of the superstition of control (19) and subjective bias (37). Therefore, the extent to which the activations produced by the illusion of control reflect the efforts of the brain to learn and to identify and filter relevant information can be isolated only in placebo-controlled studies. In summary, activation in the ventral striatum observed when participants are instructed to use their brain activation to control some external display should not be trusted as the sole index of genuine neurofeedback learning and neurofeedback reward processing as suggested by others (8) unless one can convincingly show that the pure neurofeedback related effects are stronger than those produced by sham neurofeedback. Despite the large interest in the technique of real-time fMRI neurofeedback and the large number of publications, only few placebo-controlled studies exist (30, 38). Placebo-controlled designs should be employed more frequently in the fMRI neurofeedback literature.

## Acknowledgements

We appreciate the financial support provided by the Austrian Science Fund (FWF) through grant KLI 639 and I-4184 both supporting FF. We are thankful to all participants of the study for their participation and patience.

